# High Purity Differential Expression of Genes across Spatial Domains

**DOI:** 10.64898/2026.04.20.719761

**Authors:** Bingbo Wang, Chenxi Guo, Jiaojiao He, Lin Gao

## Abstract

Spatial transcriptomics maps gene expression within tissue architecture, offering deep insights into cellular function. A key yet overlooked phenomenon is the systematic imbalance in gene regulation within spatial domains, contrasting with traditional analyses. We define and validate this as a prevalent hallmark—High-Purity Differential Expression (HiP-DEP)—where genes in a domain shift predominantly in one direction (i.e., predominantly up- or downregulated). We introduce a Spatial Purity index to quantify it. Analysis of 190 diverse datasets across diverse tissues, diseases, and technologies revealed that spatial domains possess significantly higher purity than conventional bulk (***p* = 5.4 *×* 10**^***−*11**^) or single-cell data (***p* = 6.9 *×* 10**^***−*14**^), establishing HiP- DEP as a core hallmark of spatial biology. Using our HiP-DEP framework, we show its utility in: decoding core oncogenic regulation and its spatial gradient in breast cancer; uncovering hidden phenotypic divergence between morphologically similar Alzheimer’s plaques; and identifying active cellular communication niches in normal brain tissue. By shifting focus from individual genes to coordinated transcriptional programs, HiP-DEP provides a new paradigm for precision spatial omics analysis.

## Introduction

Spatial transcriptomics (ST) (Williams et al. 2022) has rapidly advanced, revolutionizing our understanding of tissue biology by mapping gene expression within its native spatial context. It enables the analysis of complex tissue architectures and cellular interactions, and the growing generation of data from diverse systems—from tumors to developing organs—provides unprecedented resources for studying tissue organization across physiological and pathological conditions (Song et al. 2025; Schaffer et al. 2025).

A foundational step in ST analysis is identifying spatial domains, defined as contiguous tissue regions that share coherent gene expression patterns, to decipher tissue architecture and function. Determining their biological roles fundamentally involves identifying differentially expressed genes (DEGs) across them. This analysis reveals region-specific transcriptional activity, crucial for understanding local biological functions, structural specializations, and their alterations in disease, thereby linking gene activity to underlying cellular processes during progression. Currently, methods for identifying such spatial expression differences fall into three groups: (1) applying differential expression (DE) tools from bulk RNA-seq; (2) employing methods designed for single-cell RNA-seq (scRNA-seq); and (3) using algorithms specifically created for ST.

For bulk RNA-seq, expression profiles from diseased and healthy samples are compared.A common initial approach adapts established methods this domain, such as DESeq2 (Zhao et al. 2022) and Limma (Ritchie et al. 2015). DESeq2 employs a negative binomial model for rigorous statistical testing, while Limma uses a linear modeling framework with empirical Bayes moderation. However, their direct application to ST data is non-trivial, as the scale, sparsity, and spatial dependency of ST datasets necessitate specific adjustments.

For single-cell RNA-seq (scRNA-seq), a given cell type is compared against all others. Given shared characteristics like high dimensionality and sparsity, specialized methods have been developed for DE analysis. Tools like DEsingle (Miao et al. 2018) model zero-inflated counts, enabling DE detection across cell types, albeit with computational cost, as used in studies of early neuronal development (Alamin et al. 2024). Method based on Markov random field (Li et al. 2021) attempt to integrate structural relationships, though complexity often restricts use. Popular frameworks like Seurat (Sun et al. 2019; Satija et al. 2015) also provide DE workflows to identify specific markers of cell-types.

ST analysis focuses on identifying molecular features that vary across tissue architecture. A core task is the detection of Spatially Variable Genes (SVGs), which typically involves comparing gene expression in a defined Region of Interest (ROI) or spatial domain to that of the remaining tissue area. Spatially-specific algorithms are more principled. When prior domain knowledge is available, supervised methods like C-SIDE (Cable et al. 2022) can be applied, which incorporates cell type density to model expression changes in mouse brain interaction regions (Wan et al. 2023). In the absence of pre-defined domains, unsupervised detection of SVGs is employed. These can be broadly categorized by computational foundation. Statistical modelbased approaches, such as Trendsceek (Edsgärd et al. 2018), SpatialDE (Svensson et al. 2018), and SPARK (Sun et al. 2020), provide a framework for quantifying spatial heterogeneity, though computational cost can be a limitation. SPARK-X (Zhu et al. 2021) addresses this with a non-parametric strategy. Once SVGs are identified, DE analysis across spatial domains can be performed, thereby yielding a set of domain-specific DEGs characterized by significant spatial variability in expression. Tools like MERINGUE (Miller et al. 2021) utilize spatial autocorrelation statistics. Machine learning-based approaches, including spVC (Yu and Li 2024), SpaGCN (Huet al. 2021), and spaVAE (Tian et al. 2024), have been developed to capture complex spatial patterns. STAMarker (Zhang et al. 2023) is an integrated framework for domain-specific SVG identification.

Since the inception of bulk RNA-seq, DE analysis has been a cornerstone for identifying condition-specific genes, with a well-established characteristic being the relative balance between up- and down-regulated genes, visually apparent in bilaterally symmetric volcano plots (Ritchie et al. 2015). In stark contrast, our analysis of ST data reveals a fundamentally different pattern: differential expression within spatial domains frequently exhibits a pronounced imbalance, with genes strongly biased towards either up- or down-regulation, resulting in a unilateral, asymmetric volcano plot.

This observed systematic imbalance in ST, a pattern distinct from traditional analyses, raises key questions: (1) Is this a fundamental, overlooked hallmark of spatial biology? (2) How can it be defined and quantified? (3) What is its functional relevance? To answer these, we introduce the High-Purity Differential Expression (HiP-DEP) pattern. We define a Spatial Purity index (SPi) to quantify it. Using a multi-omics benchmark, we demonstrate that high-purity patterns (SPi *>* 0.7) are a prevalent and distinctive feature of spatial transcriptomics, establishing HiP-DEP as a new core principle. We then apply the HiP-DEP framework to decode its biological significance, revealing: a core oncogenic gradient in breast cancer (SPi *>* 0.97), hidden phenotypic divergence in Alzheimer’s disease plaques, and the alignment of high-purity domains with active cellular communication niches in the brain. This establishes HiP-DEP as a new paradigm for spatial omics analysis.

## Results

### From Concept to Metric: The HiP-DEP Framework

Differential expression analysis is a robust method for identifying gene sets with significant expression differences between experimental conditions or groups. We apply these techniques to characterize the expression features of specific spatial domains. A critical prerequisite for this analysis is a priori definition of spatial domains (details in Methods), which provides the necessary prior knowledge for comparing expression differences between regions.

Once domains are defined, the workflow proceeds by comparing gene expression levels between a target spatial domain and all other domains within the same tissue section to identify marker genes that define the region’s expression profile. This process typically involves identifying differentially expressed genes (DEGs) based on statistical thresholds for fold-change (FC) and adjusted p-value (p.adj).

Visualizing these results reveals a critical insight. Volcano plots for bulk and scRNA-seq data typically show a symmetrical distribution of up- and down-regulated genes (UR-DEGs and DR-DEGs) (Fig. 1). In stark contrast, ST data frequently reveals a pronounced imbalance within a given spatial domain, referred to as Region of Interest (ROI), where DEGs are overwhelmingly skewed toward either up- or down-regulation. We define this recurrent, asymmetric pattern as the High-Purity Differential Expression (HiP-DEP) pattern.

**Fig. 1.**
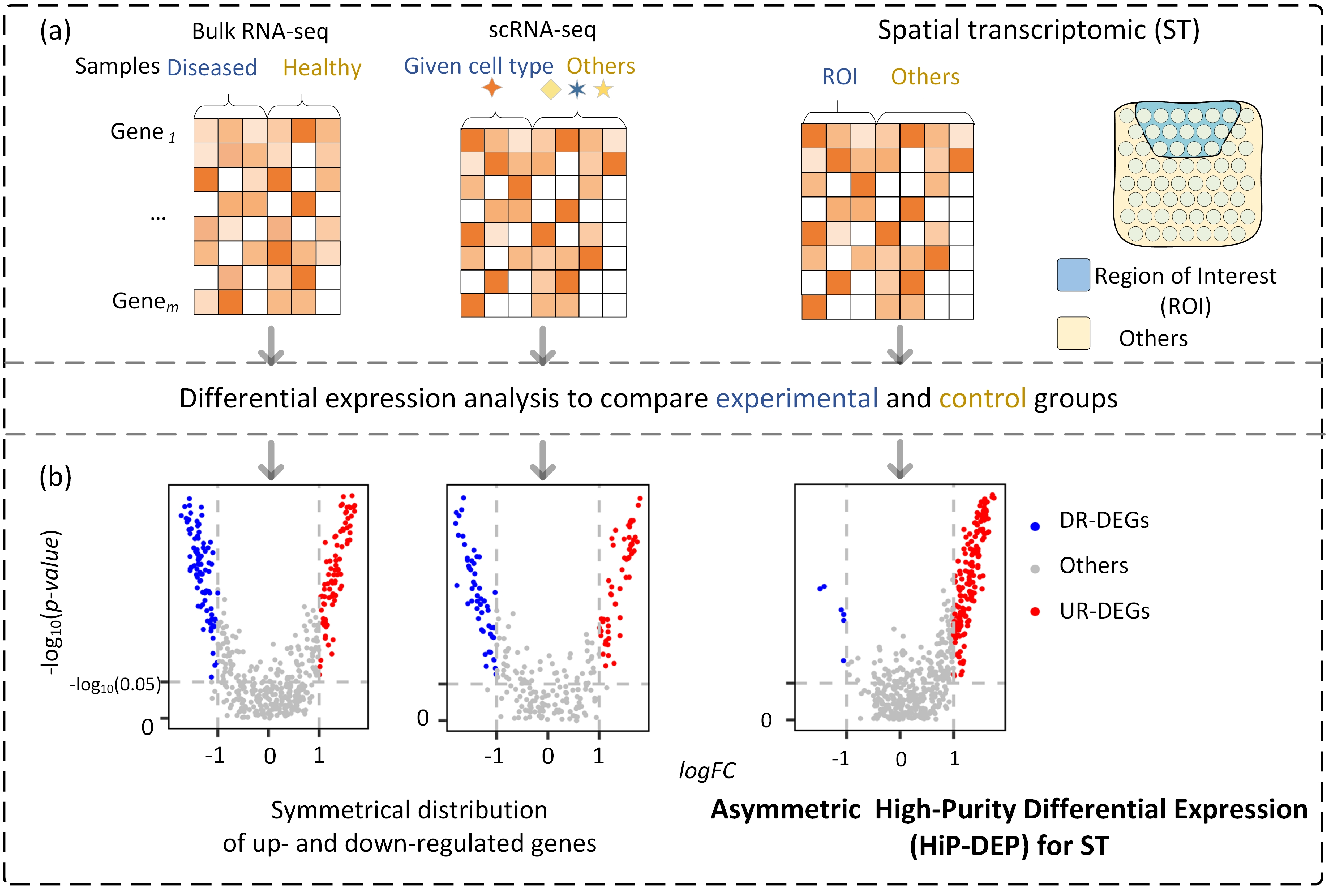
Observation of HiP-DEP. (a) Schematic of differentially expressed genes (DEGs) analysis frameworks. For Bulk RNA-seq, expression profiles from diseased and healthy samples are compared. For Single-cell RNA-seq (scRNA-seq), a given cell type is compared against all others. For Spatial Transcriptomics (ST), a Region of Interest (ROI) or spatial domain is compared to the remaining tissue area. (b) Volcano plots of DEG results. The distribution of significant DEGs (adjusted *p*-value *<* 0.05, shown as *−* log_10_(*p*-value) on the y-axis) reveals a key contrast.

To move beyond qualitative observation and statistically capture the HiP-DEP phenomenon, we formulate the Spatial Purity index (SPi), and classify spatial domains as Up-Regulation dominant (UR-domains) or Down-Regulation dominant (DR-domains) based on their predominant DEG set (Fig. 2a). This metric quantifies the dominant directional bias of gene regulation within a spatial domain: SPi = max(*N*^*up*^, *N*^*down*^)*/*(*N*^*up*^ + *N*^*down*^), where *N*^*up*^ and *N*^*down*^ are the counts of significantly up- and down-regulated genes. A high SPi-value (closer to 1) indicates a spatially coherent transcriptional program where changes are overwhelmingly concerted in one direction, signaling a unified functional or phenotypic state. A low SPi-value (closer to 0.5) reflects a balanced, mixed regulation typical of bulk analysis. Thus, the index provides a direct, continuous measure of the HiP-DEP pattern’s strength, enabling rigorous cross-domain and cross-study comparisons.

**Fig. 2.**
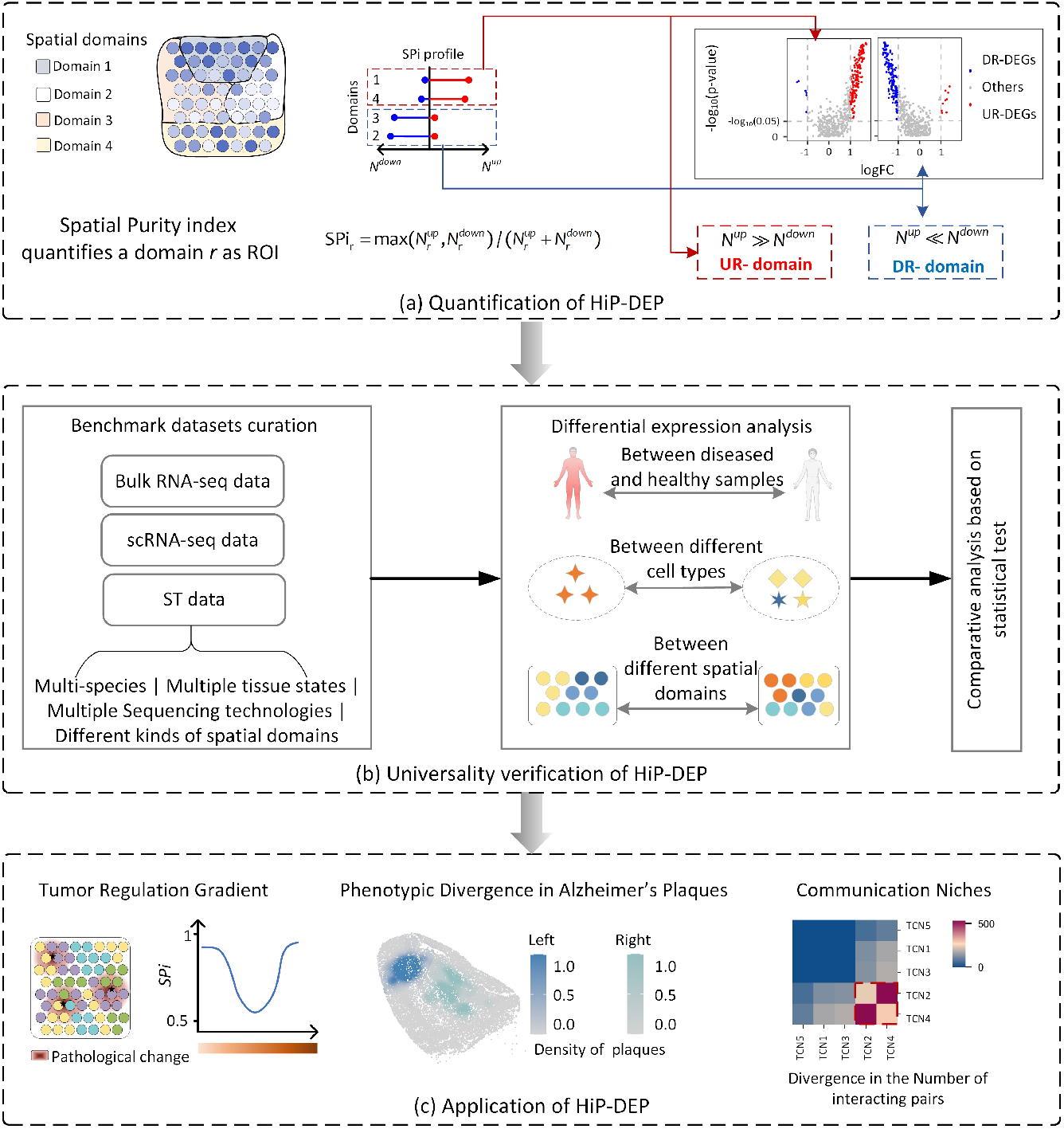
Framework of HiP-DEP Analyzer. (a) Quantification of HiP-DEP. For a spatial domain (Region of Interest, ROI), DEGs are categorized as Up-Regulated (UR-DEGs, red) or DownRegulated (DR-DEGs, blue). The unilateral bias is quantified by the Spatial Purity index (SPi). (b) Universality Verification. A diverse benchmark dataset, integrating Bulk RNA-seq, scRNA-seq, and Spatial Transcriptomics (ST) data across multiple dimensions, was established. Statistical comparison of SPi values across data types validates HiP-DEP as a hallmark of spatial data. (c) Application of HiP-DEP. The framework is applied to decode biological contexts: mapping a Tumor Regulation Gradient, revealing Phenotypic Divergence in Alzheimer’s plaques, and identifying active Communication Niches in tissue neighborhoods.

To verify HiP-DEP as a universal hallmark of spatial biology, we constructed integrated benchmark datasets containing bulk RNA-seq, scRNA-seq, and ST data. The ST benchmark is comprehensive, spanning multiple species, tissue conditions (diseased/healthy), and technologies (e.g., 10x Visium, Slide-seq v2) (Fig. 2b). Using multiple DE methods to ensure robustness, subsequent statistical validation confirmed the high prevalence and specificity of HiP-DEP across spatial domains.

Finally, we apply the HiP-DEP framework in three key contexts: within disease spatial domains to link expression patterns to pathological mechanisms, including tumor regulation gradients in breast cancer and phenotypic divergence in Alzheimer’s plaques, and within tissue cellular neighborhoods to explore their association with cell-cell communication. This end-to-end framework establishes a new paradigm for analyzing spatial transcriptomics.

Having established the HiP-DEP concept and framework, we next demonstrate its universal validity and its utility in decoding biological function. We first provide statistical evidence that HiP-DEP is a prevalent hallmark of spatial biology. We then apply the framework in three diverse contexts: to elucidate core functional regulation in breast cancer, reveal hidden phenotypic heterogeneity in Alzheimer’s disease, and identify active communication niches in normal brain tissue.

### Prevalence and specificity of HiP-DEP

To validate the universality of HiP-DEP, we established a comprehensive benchmark dataset comprising 190 spatial transcriptomic (ST) datasets, bulk RNA-seq for 18 types of cancer, and scRNA-seq for multiple cell types from 23 human tissues (details in Additional File 1 Note 1). The ST cohort encompasses multiple species, diverse tissue types, varying pathological statuses, and a range of technological platforms (Fig. 3a). The inclusion of bulk and scRNA-seq data enables a direct comparison to test the specificity of HiP-DEP to spatially resolved data.

**Fig. 3.**
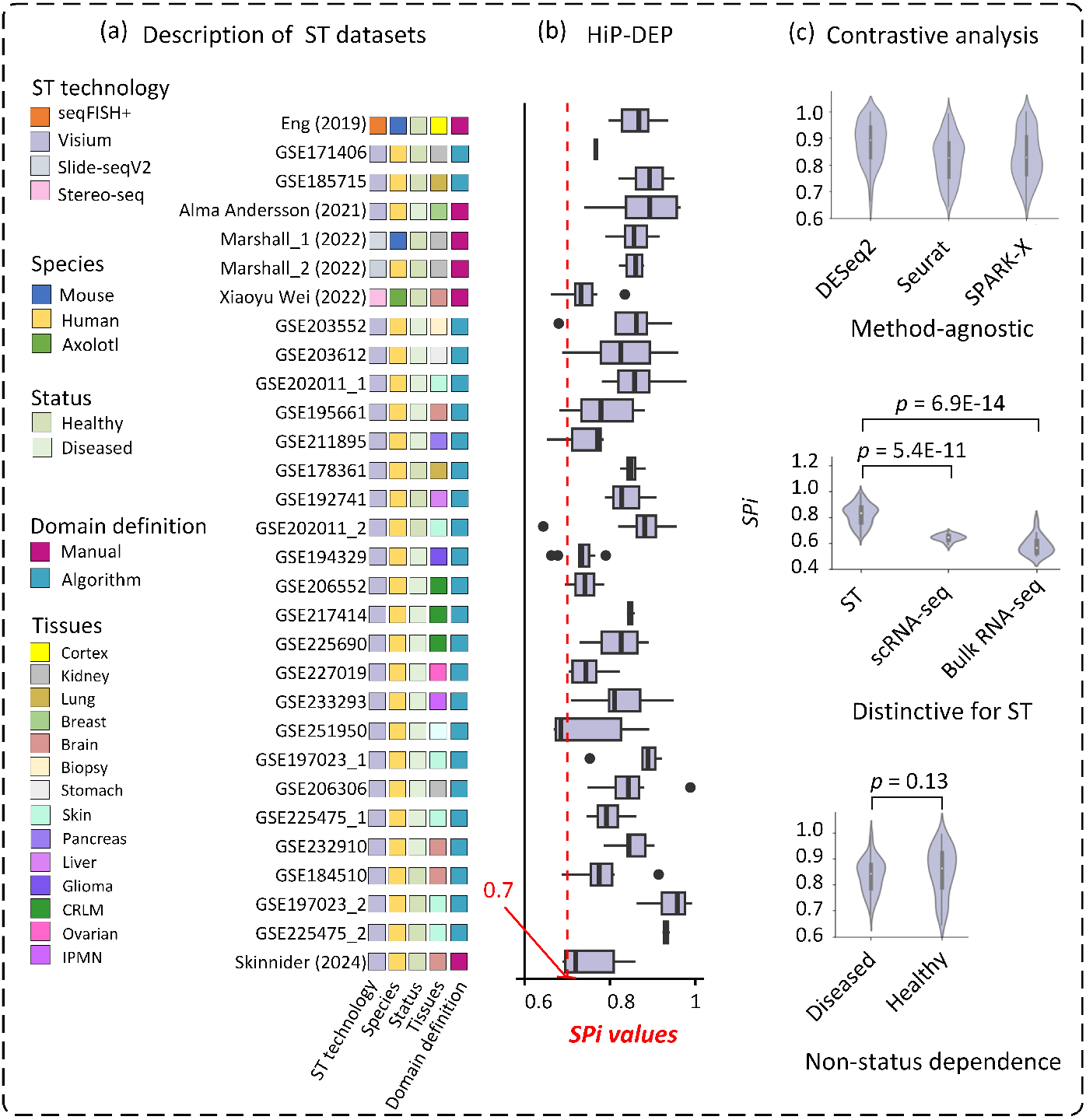
HiP-DEP across multi-dataset benchmark. (a) 190 benchmark ST datasets from 30 sources are categorized across multiple dimensions to ensure comprehensiveness: ST Technology, Species, Tissue Status (Healthy/Diseased), Domain Definition Method (Manual/Algorithmic), and Tissue of Origin. A unique identifier is listed for each dataset. (b) The plots show the SPi value for every analyzed spatial domain. The red dashed line indicates the SPi = 0.7 threshold. The distribution is significantly skewed towards high purity (SPi *>* 0.7), establishing the HiP-DEP pattern as a prevalent feature of the ST benchmark. (c) Three key comparative analyses are shown: SPi values are consistent across different differential expression analysis tools (e.g., DESeq2, Seurat, SPARK-X). The SPi of ST domains is significantly higher than that of tests from bulk or single-cell RNA-seq data. High SPi values are prevalent in both healthy and diseased tissues.

To ensure robustness, we applied multiple differential expression methods originally designed for distinct data types and bioinformatics applications: DESeq2 for bulk RNA-seq, Seurat for scRNA-seq, and SPARK-X for ST. Here, we repurposed these methods to test for differential expression between spatial domains. Fig. 3b presents the SPi values derived from SPARK-X, while Fig. 3c includes results from all three methods, demonstrating consistent trends and supporting the method-agnostic nature of our conclusions.

Analysis of the SPi revealed a marked contrast between data types. The SPi for ST data was consistently high, exceeding 0.7 across datasets (details in AdditionalFile 2 Table 1). In contrast, both scRNA-seq and bulk RNA-seq data showed significantly lower purity, with average SPi values of 0.64 and 0.58, respectively. Statistical analysis revealed a significant overall difference in SPi across the three data modalities (Kruskal-Wallis test, *p* = 2.2*×*10^*−*16^). Subsequent post-hoc comparisons demonstrated that the SPi of ST data was significantly higher than that of both bulk RNA-seq (Mann-Whitney U test, *p* = 5.4 *×* 10^*−*11^) and scRNA-seq data (Mann-Whitney U test,*p* = 6.9 *×* 10^*−*14^).

The prevalence of HiP-DEP is consistent across tissues. We found no significant difference in SPi between datasets from diseased tissues and those from healthy tissues, with both groups exhibiting consistently high SPi values (*>* 0.7). This demonstrates that HiP-DEP is a fundamental and universal feature of ST, independent of tissue pathology.

Taken together, our multi-faceted validation establishes the HiP-DEP pattern as a prevalent, specific, robust, and fundamental hallmark of ST. This conclusion is supported by the convergence of evidence from: a large and diverse benchmark dataset; the application of multiple analytical methods; rigorous statistical comparison to non-spatial data; and controlling for the variable of tissue pathology.

### Functional Regulation in Breast Cancer

To explore the link between HiP-DEP and disease biology, we applied the framework to ST data from a *HER2* -positive breast cancer tissue section (10x Visium) (Andersson et al. 2021). Pathologists annotated the tissue into seven spatial domains based on hematoxylin and eosin (HE) images: Adipose tissue, Breast glands, Cancer in situ, Connective tissue, Immune infiltrate, Invasive cancer, and Undetermined sites (Fig. 4a). Preprocessing yielded expression profiles for 11,972 genes across 613 spatial spots (details in Methods).

**Fig. 4.**
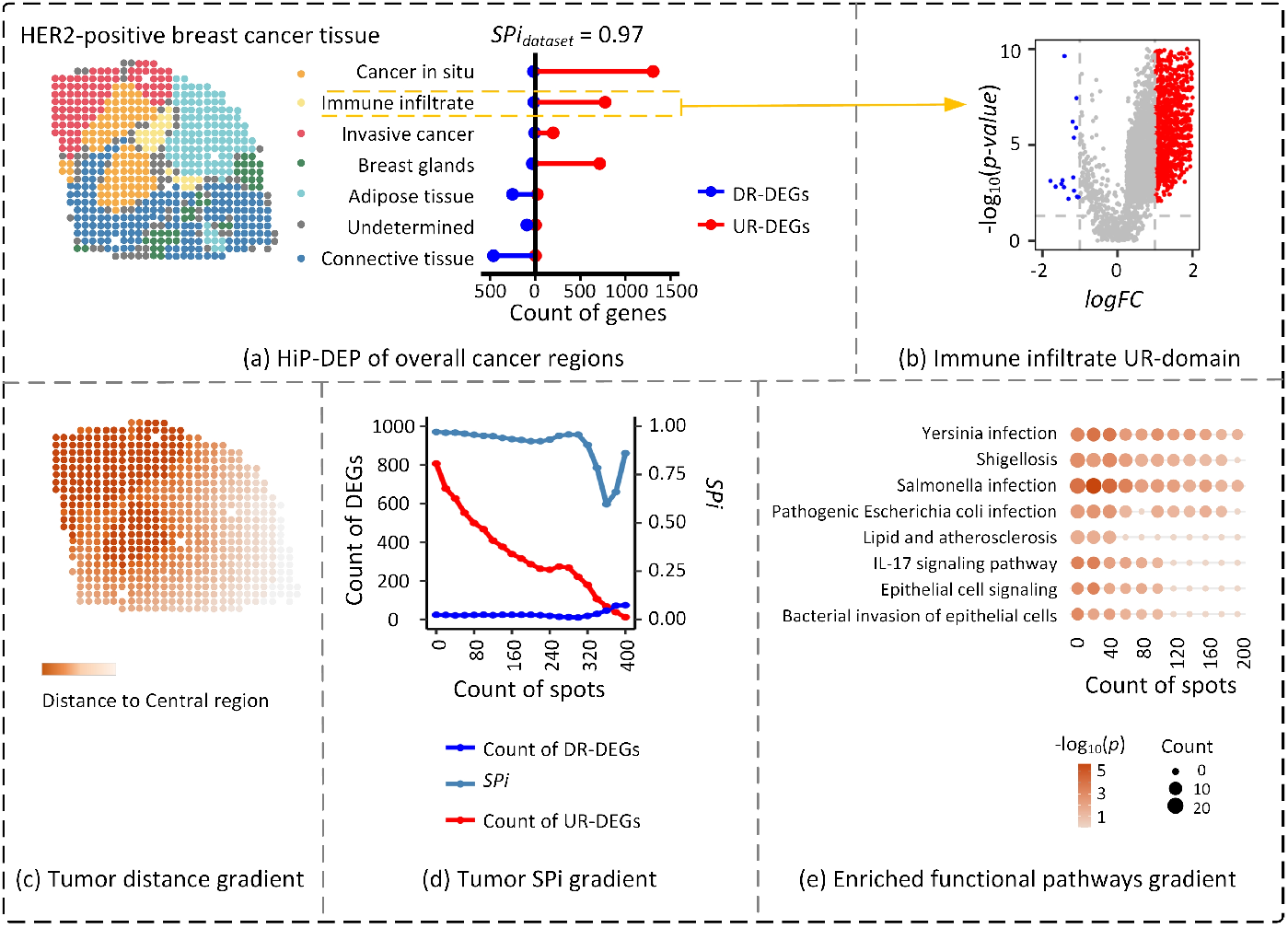
Tumor Regulation Gradient in Breast Cancer. (a) Histologically annotated spatial domains (e.g., Invasive cancer, Cancer in situ) with UR-DEG/DR-DEG counts. The dataset shows high purity (SPi = 0.97). (b) Volcano plot for the Immune infiltrate domain, illustrating the characteristic unilateral bias of HiP-DEP. (c) Spatial gradient of Euclidean distance from the tumor core region. (d) Dynamic change in UR-DEG/DR-DEG counts and SPi as analysis expands from the tumor core, showing the decay of transcriptional influence with distance. (e) Enrichment of key KEGG pathways (e.g., Yersinia infection) in the tumor core; dot color and size fade with distance, correlating with the SPi decay in (d).

Differential expression analysis of the Cancer in situ domain (compared to all other spots) revealed an extreme HiP-DEP. The SPi values from three differential expression methods were 0.99, 0.97, and 0.98. The volcano plot (SPARK-X method, Fig. 4b) showed marked asymmetry: of 1,326 differentially expressed genes (DEGs), 1,307 were up-regulated (UR-DEGs) and only 19 were down-regulated (DR-DEGs), definitively classifying it as an Up-Regulation-dominant (UR) domain. Global analysis across all domains confirmed widespread high purity, with overall SPi values of 0.91, 0.88, and 0.97. Cancer in situ, Immune infiltrate, Invasive cancer, and Breast glands were UR domains, while Adipose tissue and Connective tissue were Down-Regulation-dominant (DR) domains (Fig. 4b).

We next investigated how the disease core (Cancer in situ and Invasive cancer) influences its microenvironment. We iteratively expanded this core by adding the spatially nearest neighboring spots and re-calculated differential expression (Fig. 4c). The pattern shifted dynamically: initial expansion reduced UR-DEGs while DR-DEGs stayed stable; after adding 300 spots, DR-DEGs began to increase. The SPi declined gradually until 320 spots were added, then dropped sharply, reaching its lowest point at 360 spots. This indicates a spatial gradient: proximal regions maintain high expression (UR domain), but the core’s influence wanes with distance, leading to decreased expression and a transition to a DR domain (Fig. 4d).

KEGG enrichment analysis of these up-regulated genes in the core identified the top pathways associated with breast cancer (*p <* 0.05), including Yersinia infection (immune evasion) and IL-17 signaling (inflammation/proliferation) (Fig. 4e). The significance of these pathways decayed with distance. For example, expanding the region by 200 spots increased the p-value for the Yersinia infection pathway from 6 *×* 10^*−*4^ to 0.04, and for the IL-17 signaling pathway from 3.8 *×* 10^*−*4^ to 1. This attenuation in pathway significance correlated with the decrease in SPi, suggesting the disease core spatially modulates its microenvironment via key oncogenic pathways, with effects diminishing over distance.

### Phenotypic Heterogeneity in Alzheimer’s Disease

Spatial transcriptomics enables the analysis of gene expression across continuous tissue gradients, such as those defined by pathological features. We applied HiP-DEP to Slide-seq V2 data from the hippocampus of an Alzheimer’s disease (AD) mouse model (Zhou et al. 2020), where the pathological feature is *β*-amyloid (A*β*) plaque density. The goal was to examine the relationship between this continuous pathological measure and differential expression patterns. The raw dataset contained 27,222 genes across 15,092 spatial spots. After preprocessing (filtering genes with expression in *<* 0.1% of spots and spots with expression in *<* 0.5% of genes), 13,031 genes and 4,108 spots were retained for analysis.

We focused on two core pathological regions with plaque density *≥* 1, designated as the left plaque and right plaque (Fig. 5a). Differential expression analysis (SPARK-X method) of each plaque versus all other spots revealed a high-purity pattern (overall dataset SPi = 0.79). However, the two plaques exhibited divergent directional biases. The Right plaque was an UR domain (SPi = 0.90), while the Left plaque was a DR domain (SPi = 0.85) (Fig. 5b), indicating that spatial domains with identical pathology can harbor distinct transcriptional states.

**Fig. 5.**
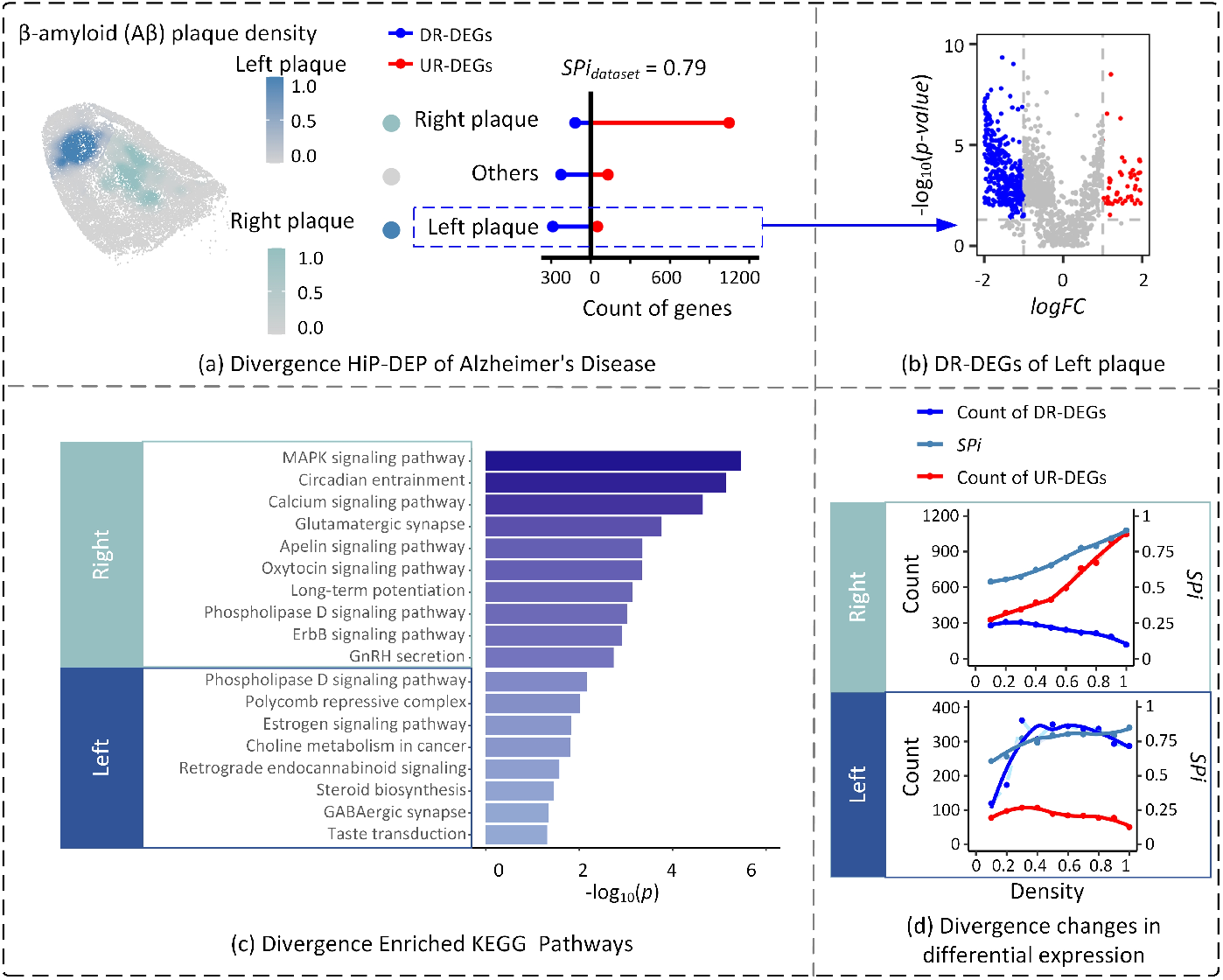
Phenotypic Divergence in Alzheimer’s Plaques. (a) Two adjacent A*β* plaques exhibit divergent HiP-DEP patterns: the Right plaque is a UR-domain (SPi = 0.90), the Left plaque is a DR-domain (SPi = 0.85). (b) Volcano plot for the DR-domain (Left plaque). (c) Distinct KEGG pathways are down-regulated in the DR-domain (e.g., Estrogen signaling) and up-regulated in the UR-domain (e.g., MAPK signaling), indicating opposing functional states. (d) The strength of the SPi for each plaque decreases as the analyzed region expands into areas of lower plaque density, linking HiP-DEP to core pathology.

KEGG enrichment analysis of the UR-DEGs in the Right plaque and the DR-DEGs in the Left plaque revealed functionally opposing microenvironments (Fig. 5c). The Right plaque (UR domain) was enriched for pathways promoting neuroinflammation and excitotoxicity (e.g., MAPK signaling, Calcium signaling). In contrast, the Left plaque (DR domain) showed downregulation of pathways critical for metabolic homeostasis, epigenetic regulation, and lysosomal clearance (e.g., Polycomb repressive complex, Estrogen signaling, Phospholipase D signaling). Notably, the Phospholipase D pathway was differentially regulated: *Pld1* was up-regulated in the Right plaque (potentiating A*β* deposition), while *Pld3* was down-regulated in the Left plaque (impeding protein clearance). This suggests a synergistic deleterious effect where one plaque promotes inflammatory damage and the other exhibits a collapse of cellular defense mechanisms.

We further investigated how plaque density correlates with the HiP-DEP pattern. Analyzing domains of varying density revealed that SPi in both plaques decreased as plaque density decreased (Fig. 5d), but the underlying regulatory shifts were distinct. In the left plaque (DR-domain), lower density was associated with a progression from a Down-Regulation-dominant state towards a balanced mix of UR-DEGs and DR-DEGs. Conversely, in the Right plaque (UR-domain), lower density shifted the pattern from an Up-Regulation-dominant state towards a balanced state. This divergence demonstrates that plaque density is a critical pathological feature in Alzheimer’s disease, with the strength and direction of the high-purity pattern being quantitatively linked to local pathological burden.

In summary, HiP-DEP analysis of AD tissue uncovered hidden phenotypic heterogeneity: two histologically similar A*β* plaques represent functionally distinct microenvironments—one characterized by active inflammatory signaling and the other by impaired homeostatic defense. The correlation between plaque density and SPi further establishes plaque density as a critical continuous variable shaping local transcriptional polarity in AD.

### Communication Niches in Tissue Neighborhoods

Tissue cellular neighborhoods (TCNs) (Hu et al. 2024) are spatially defined multicellular units critical for tissue organization and intercellular communication. To investigate whether HiP-DEP patterns correspond to functional niches, we analyzed a 10x Visium spatial transcriptomics (ST) dataset of the human dorsolateral prefrontal cortex (DLPFC) (Maynard et al. 2021). We focused on sample DLPFC-151670, which contained expression data for 33,538 genes across 4,992 spatial spots. The tissue was first annotated into five distinct TCNs using CytoCommunity (Fig. 6a). To ensure analysis robustness, genes expressed in fewer than 0.1% of spots were filtered out, resulting in the removal of 16,142 low-quality genes.

**Fig. 6.**
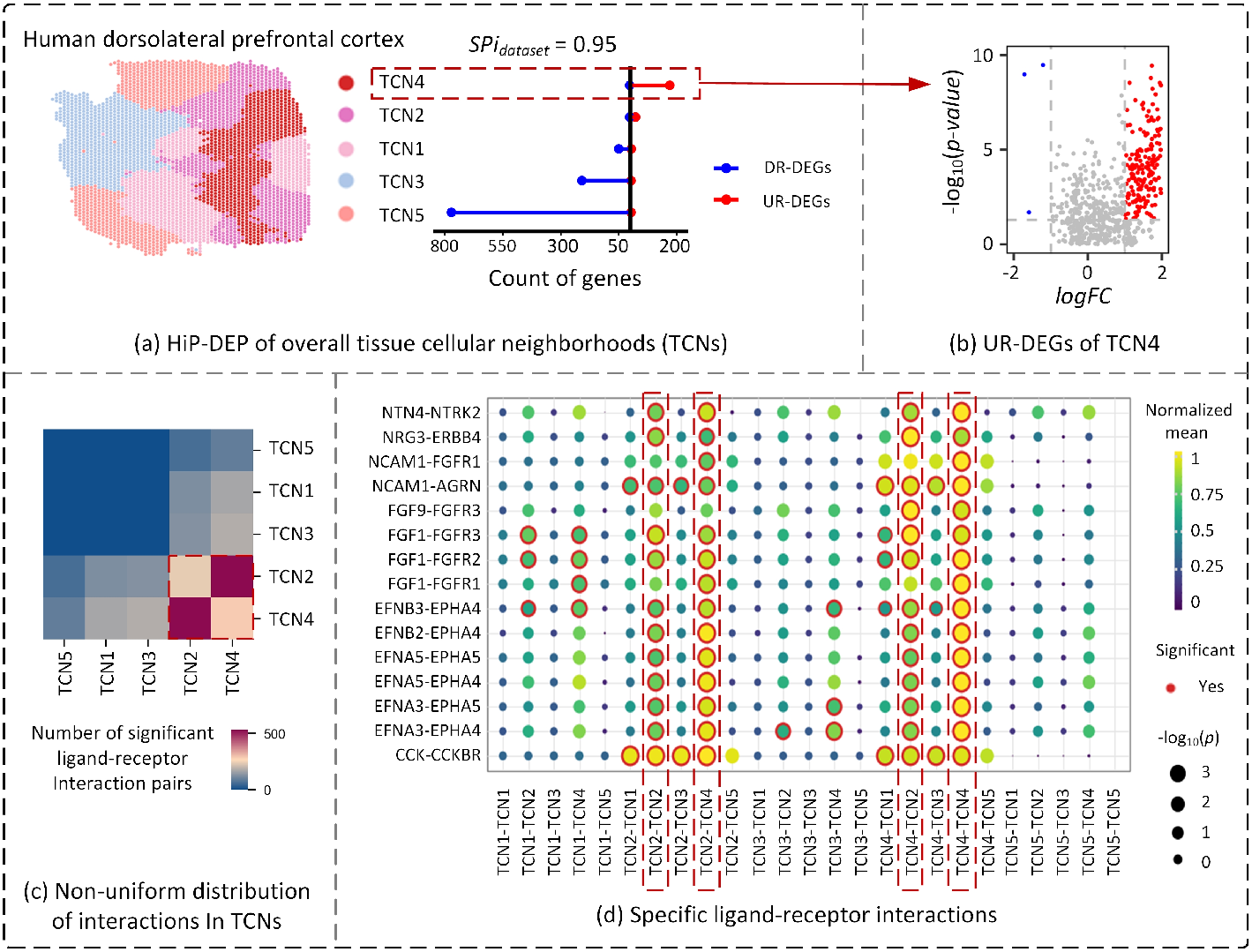
Communication Niches in Tissue Neighborhoods. (a) Bar plot shows the number of up- and down-regulated differentially expressed genes (UR-DEGs and DR-DEGs) for each TCN (TCN1–TCN5) in the human dorsolateral prefrontal cortex. The overall SPi = 0.95 indicates a tissue-wide high-purity pattern. (b) Volcano plot confirms TCN4 as a high-purity UR-domain, with genes significantly up-regulated compared to other tissue regions. (c) A heatmap visualizes the number of significant ligand-receptor interactions (*p <* 0.05) between each pair of TCNs. The strongest interactions are concentrated within the high-purity UR-domains, particularly TCN4 and between TCN2 and TCN4. (d) Enriched interaction pairs within the active TCNs are detailed. Color represents the normalized mean expression level, and red circles indicate the significant ligand-receptor interactions.

Applying HiP-DEP to each TCN revealed a striking, tissue-wide high-purity pattern (overall Spatial Purity index, SPi = 0.95) (Fig. 6a). TCN4 and TCN2 were identified as strong Up-Regulation (UR) domains, while TCN1, TCN3, and TCN5 were classified as Down-Regulation (DR) domains.

We hypothesized that the identified UR domains might represent hotspots of active cellular crosstalk. To test this, we performed a ligand-receptor interaction analysis across TCNs using CellPhoneDB (Troulé et al. 2025). The analysis revealed a nonuniform distribution of significant interactions (*p <* 0.05) (Fig. 6c). Strikingly, the two UR domains, TCN2 and TCN4, exhibited the highest number of significant ligand-receptor pairs, both internally and between each other, indicating they are the most active communication niches.

A focused analysis of the significant interactions specific to TCN2 and TCN4 highlighted key signaling pathways (Fig. 6d). These included interactions between EFNA-class ligands and EPHA receptor (Klein 2012; Cramer and Miko 2016), which are known to activate MAPK, Ras, and Rap1 pathways involved in neural development and plasticity. The co-occurrence of high-purity upregulation and enriched ligand-receptor activity in TCN2 and TCN4 suggests that their coordinated, elevated gene expression underpins a state of heightened cellular communication and metabolic activity.

In summary, applying HiP-DEP to the architecture of normal brain tissue successfully pinpointed specific TCNs as high-purity UR domains. These transcriptional hotspots spatially coincided with the tissue’s most active cell-communication niches, demonstrating that the HiP-DEP framework can identify and functionally characterize discrete units of cellular crosstalk within a complex tissue microenvironment.

## Discussion

The advent of spatial transcriptomics (ST) has enabled the mapping of gene expression within intact tissue architecture, revealing cellular interactions in their native context. A critical task in ST analysis involves identifying spatial domains—regions of coherent expression—and performing differential expression (DE) analysis to decode their functional identity. However, existing analytical frameworks have largely overlooked a systematic feature we consistently observe: a pronounced imbalance between up- and down-regulated genes within spatial domains, manifesting as a unilateral asymmetry in DE volcano plots, distinct from the balanced patterns typical of bulk or single-cell RNA-seq. We termed this phenomenon the High-Purity Differential Expression (HiP-DEP) pattern.

To investigate HiP-DEP, we introduced a Spatial Purity index (SPi) for quantification and developed a comprehensive analytical framework. Using a set of benchmark datasets spanning multiple species, tissues (healthy and diseased), and technologies, we statistically established HiP-DEP as a prevalent and specific hallmark of spatial biology. The purity of differential expression in spatial domains significantly exceeded that in bulk or single-cell data.

Applying the HiP-DEP framework, we uncovered its utility in decoding distinct biological processes. In breast cancer, high-purity patterns delineated core tumor regions and revealed a spatial gradient of regulatory influence on the microenvironment. In Alzheimer’s disease, HiP-DEP exposed hidden phenotypic heterogeneity between morphologically similar amyloid plaques, linking differential plaque density to divergent transcriptional states. In normal brain tissue, high-purity Up-Regulation-dominant (UR) domains spatially coincided with the most active cell-communication niches identified by ligand-receptor interaction analysis.

The determinants of HiP-DEP appear linked to the mean expression level of a spatial domain relative to its surroundings. A domain with a significantly different average expression profile tends to exhibit high-purity differential expression. Beyond global expression levels, future studies using single-cell or subcellular resolution ST could elucidate how specific cell types, states, and intercellular regulatory networks contribute to these coordinated expression patterns. Integrating multi-omic data layers may further reveal the upstream biological mechanisms.

In summary, HiP-DEP provides a novel paradigm for analyzing spatial transcriptomics by shifting focus from individual variable genes to the systematic directional bias of transcriptional programs within tissue neighborhoods. It serves as a powerful framework for identifying functionally coherent spatial units, revealing hidden heterogeneity, and pinpointing active communication niches, thereby advancing precision spatial omics.

## Methods

The following sections detail the computational and statistical methods underpinning the HiP-DEP framework. We begin by describing the data curation and preprocessing steps applied to all data types. Next, we outline the procedures for defining spatial domains in ST data and corresponding pseudo-domains in non-spatial data to enable a fair comparative analysis. We then specify the differential expression analysis pipelines, detail the calculation of the Spatial Purity index (SPi), and present the statistical testing used to establish the universality and specificity of the high-purity pattern.

### Data Curation and Preprocessing

Spatial transcriptomics (ST) data are represented as an expression count matrix 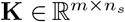, where rows correspond to *m* genes (gene set *G* = *{g*_1_, *g*_2_, …, *g*_*m*_*}*) and columns correspond to *n* spatial spots (spot set 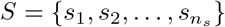. The spatial coordinates are stored in a matrix 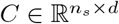, where *d* denotes the spatial dimensions.

To mitigate the impact of low-quality data and high dropout rates characteristic of ST technologies, we applied a standardized preprocessing pipeline (Additional File 1 Note 2, Algorithm 1). First, we calculated the non-zero expression ratios for each gene (Gratio = 0.1%) and each spot (Sratio = 0.5%) from the raw count matrix **K**. Genes and spots with ratios below predefined thresholds were filtered out. This step removes genes with extremely low detection rates and spots with an insufficient number of detected genes, reducing technical noise.

Subsequently, the filtered expression matrix was normalized using the Transcripts Per Million (TPM) method (Zhao et al. 2021) to account for variations in sequencing depth and technical biases across spots. This normalization yields a spot-specific relative expression measure, facilitating comparative analysis across the spatial landscape.

### Spatial Domains and Pseudo-domains Definition

The definition of spatial domains is a critical prerequisite for all subsequent analyses. In this study, domains are established through two primary approaches: expert annotation and computational detection, ensuring robustness and biological relevance. Expert Annotation. When high-quality histological images are available, pathologists can manually delineate spatial domains based on distinct morphological features.

This provides a gold standard grounded in pathological expertise.

Computational Detection. For automated, data-driven domain identification, we employed two state-of-the-art algorithms to assess the consistency of the HiP-DEP pattern across different detection methods. GraphST (Long et al. 2023): A graph supervised contrastive learning method designed for general spatial domain detection. It demonstrates high accuracy across diverse tissue types and technological platforms. CytoCommunity(Hu et al. 2024): An algorithm that identifies tissue cellular neighborhoods by integrating cell type composition and spatial proximity. The neighborhoods it identifies are instrumental in uncovering condition-specific cell-cell communication patterns.

For non-spatial data (bulk and scRNA-seq), analogous pseudo-domains are generated by applying standard clustering algorithms (e.g., Louvain(Blondel et al. 2008)) directly to the expression profiles to enable comparative analysis.

### Differential expression analysis

For a given ST dataset, differential expression analysis is performed for each annotated spatial domain. Let *R* be the set of all spatial domains. For a specific domain *r ∈ R*, we define its corresponding spot set as 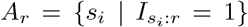, where 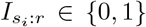 indicates whether spot *s*_*i*_ belongs to domain *r*. The gene expression matrix for domain *r* (experimental group) and for all spots outside domain *r* (control group) are extracted from the preprocessed matrix **K**. Let **k**_*i*_ denote the *i*-th column vector of matrix **K**, representing the gene expression profile of spot *s*_*i*_.

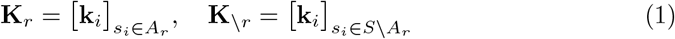

A differential expression analysis function *f*_DE_ is then applied to compare the two groups:

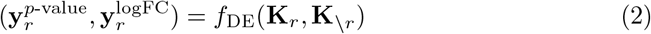

This function returns two *m*-dimensional vectors for domain 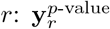 containing the adjusted *p*-values (*p*.adj) and 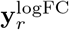 containing the log_2_ fold-changes (logFC) for all genes. The *p*-value assesses the statistical significance of expression differences, while the logFC quantifies their direction and magnitude. A positive logFC indicates up-regulation in domain *r*, whereas a negative value indicates down-regulation.

Differentially expressed genes (DEGs) are identified by applying thresholds on both statistical significance and fold-change. Genes with *p*.adj *<* 0.05 and logFC *>* 1 are classified as up-regulated DEGs (UR-DEGs), and genes with *p*.adj *<* 0.05 and logFC *< −*1 as down-regulated DEGs (DR-DEGs). The counts of UR-DEGs and DR-DEGs for domain *r* are calculated as:

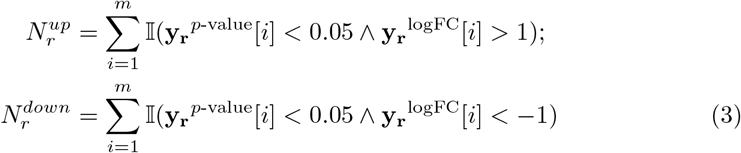

where 𝕀 (*·*)is the indicator function. This process is applied systematically to all spatial domains *r ∈ R*, as formalized in Algorithm 2 (Additional File 1 Note 2), to generate a complete set of differential expression results.

### Calculation of the Spatial Purity index (SPi)

To quantify the directional bias of differential expression within a spatial domain, we introduce the Spatial Purity index (SPi). This metric measures the degree of imbalance between up-regulated (UR-DEGs) and down-regulated (DR-DEGs) genes in a given domain, analogous to cluster purity in clustering evaluation.

For a spatial domain *r* with 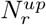 UR-DEGs and 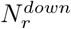 DR-DEGs, the SPi is defined as:

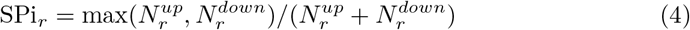

The value of SPi_*r*_ ranges from 0.5 to 1. SPi_*r*_ = 1 indicates that all DEGs in domain *r* are consistently either up-regulated or down-regulated, representing a perfect high-purity pattern. SPi_*r*_ = 0.5 indicates an equal number of UR-DEGs and DR-DEGs, reflecting a balanced, non-pure pattern typical of bulk analysis. In the extreme case where no differentially expressed genes are identified 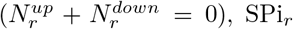 is assigned a value of 0.5.

The overall SPi for an entire ST dataset, denoted as *SPi*_*dataset*_, is calculated as the mean of the SPi values across all spatial domains:

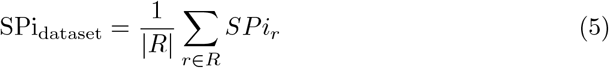

where *R* is the set of all spatial domains and |*R*| is the total number of domains. Thus, a dataset with SPi_dataset_ *>* 0.7 is considered to exhibit a pronounced highpurity pattern, whereas a value approaching 0.5 indicates a balanced, non-pure pattern typical of conventional analyses.

This metric allows systematic comparison of transcriptional bias across domains, datasets, and data types, serving as the core quantitative tool for identifying HiP-DEP.

### Statistical Testing and Validation

To statistically validate the universality and specificity of the HiP-DEP pattern, we performed comparative analyses of the Spatial Purity index (SPi) distributions across data types. First, we used the non-parametric Kruskal-Wallis test to assess whether the median SPi values differed significantly among spatial, bulk, and single-cell RNA-seq data types. Upon obtaining a significant overall result, we conducted post-hoc pairwise comparisons using the Mann-Whitney U test to determine which specific data type pairs exhibited significant differences. This two-step approach rigorously tests our hypothesis that SPi is significantly higher in spatial transcriptomics data than in nonspatial data. All p-values from pairwise tests were adjusted for multiple comparisons using the Benjamini-Hochberg method to control the false discovery rate. Detailed test results, including exact p-values and effect sizes, are reported in the Supplementary Information (Additional File 1). This statistical framework establishes that high-purity differential expression is a prevalent and distinctive feature of spatial biology.

## Data availability

Benchmark datasets (190 ST datasets, bulk RNA-seq for 18 cancer types, and scRNA-seq from 23 human tissues): NCBI GEO (Barrett et al. 2012) (https://www.ncbi.nlm.nih.gov/geo/), Xena Browser (Goldman et al. 2020) (https://xenabrowser.net/), Single Cell Portal (Tarhan et al. 2023) (https://singlecell.broadinstitute.org/), 10x Genomics (Zheng et al. 2017) (https://www.10xgenomics.com/), Zenodo(https://zenodo.org/),, Dryad (https://datadryad.org/) and SpatialOmics (Yuan et al. 2023) (https://gene.ai.tencent.com/SpatialOmics/).

The datasets and computational results supporting the three main case studies (Functional Regulation in Breast Cancer, Phenotypic Heterogeneity in Alzheimer’s Disease, and Communication Niches in Tissue Neighborhoods) are openly available in the GitHub repository at https://github.com/wangbingbo2019/HiP-DEP.

## Authors contributions

BBW: conceived and designed the experiments. JJH, CXG: performed the experiments. JJH, CXG: analyzed the data. BBW, JJH, CXG: wrote the paper. LG: revised the paper.

## Acknowledgements

We would like to thank the developers of all tools mentioned in this paper. Without the software they developed, the presented work could not exist.

## Supplementary Information

**Additional File 1**: Curation of benchmark datasets and Detailed process of method.

**Additional File 2**: Spatial Purity index results in the 190 ST datasets (Table 1).

